# The effectiveness of glass beads for plating cell cultures

**DOI:** 10.1101/241752

**Authors:** Alidivinas Prusokas, Michelle Hawkins, Conrad A. Nieduszynski, Renata Retkute

## Abstract

Cell plating, the spreading out of a liquid suspension of cells on a surface followed by colony growth, is a common laboratory procedure in microbiology. Despite this, the exact impact of its parameters on colony growth has not been extensively studied. A common protocol involves the shaking of glass beads within a petri dish containing solid growth media. We investigated the effects of multiple parameters in this protocol - the number of beads, the shape of movement, and the number of movements. Standard suspensions of *Escherichia coli* were spread while varying these parameters to assess their impact on colony growth. Results were assessed by a variety of metrics - the number of colonies, the mean distance between closest colonies, and the variability and uniformity of their spatial distribution. Finally, we devised a mathematical model of shifting billiard to explain the heterogeneities in the observed spatial patterns. Exploring the parameters that affect the most fundamental techniques in microbiology allows us to better understand their function, giving us the ability to precisely control their outputs for our exact needs.

## I. INTRODUCTION

A prominent technique in microbiology is cell plating to enable colony growth on nutrient agar. The goal of cell plating is to separate cells contained within a small sample volume by homogeneously distributing the cell suspension over the surface of a plate. This results in the formation of discrete colonies after incubation, that can be enumerated or subjected to further analysis. Counting the number of colonies is used in many cell culture protocols: detection of bacteria in food samples or clinical specimens [1]; measuring progenitor stem cell content [2]; constructing gene-knockout libraries [3]; cell-based DNA cloning [4]; testing resistance to drugs [5]; studying evolution of antibiotic resistance [6]; activation of mucosal-associated invariant T cells [7]. Successful cultivation of cells depends critically on the choice of appropriate growth media, cell plating method and incubation conditions.

A colony is defined as a visible cluster of cells growing on the surface of a medium, presumably derived from a single cell. These single progenitor cells are also called Colony Forming Units (CFUs), which provide an estimate of the number of viable cells. The more homogeneously cells are spread on the surface during plating, the better separated the resulting cell colonies, leading to a more precise CFUs estimation. It is expected that the number of colonies formed is linearly proportional to the concentration of viable cells in the suspension, provided that the suspension was mixed and spread well. If the suspension is not spread well, clusters of cells will be formed which visually resemble a single colony and therefore will be enumerated as a single colony [8]; furthermore bias may be introduced when analysing genetic variations and heredity of such clusters of cells [9]. To proceed with further procedures, the cells need to be restreaked to get single colonies which takes extra incubation time.

There are two strategies for cell plating: spread-plating with a turntable rod or spread-plating with glass beads [10, 11]. Using the first method, a small volume of a cell suspension is spread over the plate surface using a sterile bent glass rod as the spreading device. The second method, which is the subject of our study, involves shaking glass beads over the surface of the plate. This technique is also known as the Copacabana method [12, 13].

The protocol for using sterile glass beads for dispersion of cells on solid media has the following steps [11, 12]:(1) a cell suspension is dispensed onto the middle of a round Petri dish containing solid media; (2) spherical glass beads are poured into the middle of the plate; (3) the plate lid is closed; (4) the plate is agitated with a shaking motion so that the glass beads roll over the entire surface of the plate; (5) the plate is then inverted to remove the beads before incubation.

In this study we investigate how the way a plate is agitated influences spread of CFUs. We perform experiments with the bacteria E.coli and use a model from statistical mechanics - billiard, to explain heterogeneities in the observed spatial patterns. Billiard is a type of mathematical model describing a dynamical system where one or more particles moves in a container and collides with its walls [14]. Billiard systems based on the wave dynamics in cavities, acoustic resonance in water, atoms bouncing off beam of light and quantum dots have been studied over several decades both experimentally and theoretically [15–21]. We introduced a shifting billiard as a conceptual abstraction of the cell plating with glass beads. Numerical analysis of the dynamics of this simple yet efficient billiard, indicated a close relationship between plate movement and quality of colony distribution.

## II. CELL PLATING EXPERIMENT

The experiments were performed on circular plates with 88 mm diameter. Sterile 4 mm glass plating beads (manufactured by Sigma-Aldrich) were used for the dispersion of cells over the surface of a plate. This gives the ratio between glass bead and plate *d* = 4*/*88 = 0.045.

We explored following movement of a plate:

- L-shape. Plate is moved on a trajectory resembling the horizontally reflected letter “L”: up-right-left-down (↑→←↓).
- Up-down. Plate is moved up and down along the vertical axes (↑↓).
- *Hourglass. Plate is moved on a trajectory resembling the hourglass: diagonal up-left-diagonal down-right (*↗←↘.).

We called the full cycle of movement, where a plate returns to its initial position, a loop.

*E. coli* K-12 MG1655 was incubated in LB broth overnight [22]. A single *E. coli* colony was sampled from a streaked plate, placed within the LB broth, and incubated in a 37°C shaker overnight until saturation. This overnight stock was serially diluted by factors of 10 in 56/2 salts to the required dilution. 100 *µl* of the cell dilution was plated on LB agar 1.8% W/V, utilising the above motions, with a varying number of loops and beads. These plates were incubated overnight at 37 °C, and subsequently imaged. Image analysis is described in Appendix A.

## III. ASSESSING EFFECTIVENESS OF PLATING METHOD

The colony counts were performed automatically using a custom made code for image analysis, more details are given in Appendix A. We used the following metrics to assess plating methods: number of CFUs, *n*_*CFUs*_; mean distance to nearest CFU, *d*_*NB*_; and variability and uniformity in CFU spatial distribution on a plate. To determine the spatial variability, we have grouped CFUs into 5° intervals, calculated the number of CFUs in that arc, and estimated the coefficient of variation between these numbers, *CV*_*S*_. Next we have calculated distances of CFUs to the center of the plate, binned into 1*mm* intervals and calculated an average difference between observed frequencies and expected frequencies, *m*_*S*_. Expected frequencies for the later metrics were assumed to follow the triangular distribution. Larger values of *n*_*CFUs*_ and *d*_*NB*_ correspond to a better spread, while smaller values of *CV*_*S*_ and *m*_*S*_ indicate a more uniformly spread of colonies. Figure 1 shows graphical representation of these metrics, which gives the following values: *n*_*CFUs*_ = 335, *d*_*NB*_ = 2.8, *CV*_*S*_ = 0.46 and *m*_*S*_ = 2.3.

**FIG. 1.**
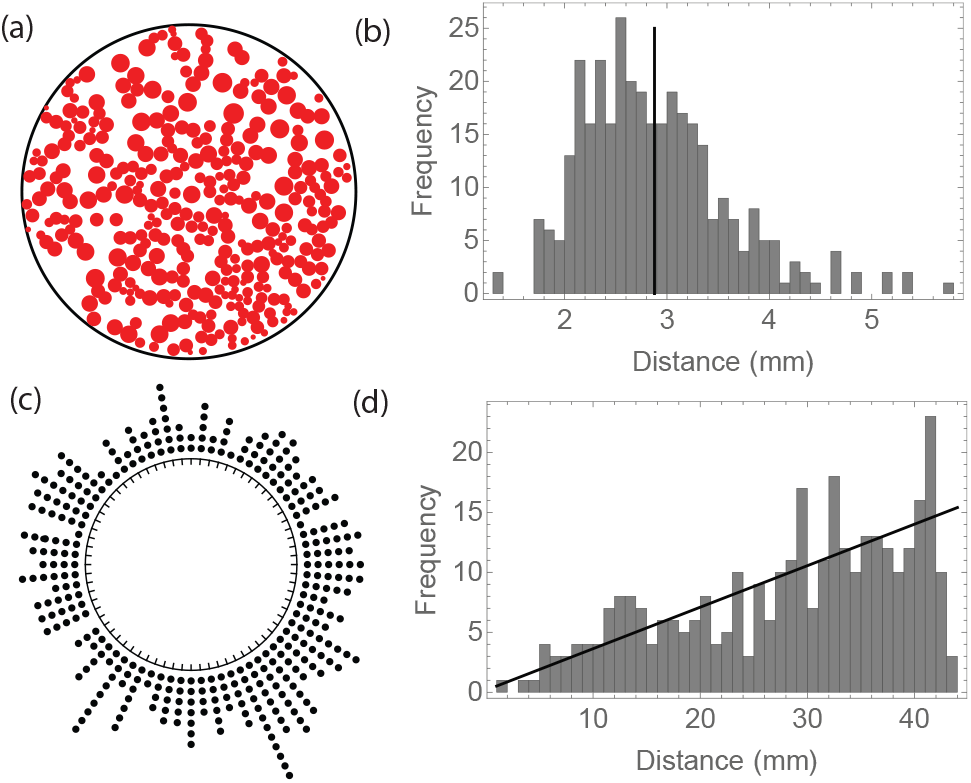
Metrics to assess the effectiveness of cell plating: (A) Counting the number of CFUs, *n*_*CFUs*_. (B) Calculating mean (dashed line) of the distribution of distances to nearest CFUs, *d*_*NB*_. (C) Dividing plate into 5° arcs, determining the number of CFUs in that arc, and calculating the coefficient of variation, *CV*_*S*_. (D) Calculating the mean difference between observed and expected distances from CFUs and plate center, *m*_*S*_; black line shows frequency based on a triangular distribution.

## IV. THE MODEL

### A. Shifting billiard

First, we start with a circular billiard which is stationary in space. Without loss of generality, we set the radius of the circle to be 1. A single particle is moving without friction between the boundary of the unit circle. Particle movement is defined by the elastic collision rule: the angle of reflection is equal to the angle of incidence [23]. The dynamics of particle movement can be described in terms of collision maps, where *θ*_*k*_ denotes the point of the collision and *φ*_*k*_ denotes the reflection angle for *k*^*th*^ collision, *k* = 1, .., *n*. We adopted a system where *θ* = 0 corresponds to a point (1, 0) in a Cartesian coordinate system. Figure 2 (a) shows a collision of a particle with the boundary at the point *θ* = *π* and 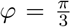. The dynamics of a particle in the circle is fully integrable and is described by the following properties: (i) *φ*_*k*_ = *φ*_1_, i.e. the angle of the reflection does not change; (ii) the coordinates of the point of collision satisfies *θ*_*k*_ = (*θ*_1_ + 2*kφ*_1_) mod 2*π*; (iii) if *φ < π* and is a rational multiple of *π*, then the particle trajectory is periodic with the particle following the sides of some regular polygon; (iv) otherwise, if 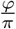 is irrational, then the particle trajectory densely fills the ring between boundary of the unit circle and the boundary of a smaller circle with radius cos^2^(*φ*) [14]. For the case (iii), the number of polygon sides depends on the reflection angle. For example, if 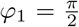, the polygon is a square and particle moves in period-4 orbit along sides of the square; and for *φ*_1_ = *π* the particle runs forth and back along a diameter of the circle and is locked in period-2 orbit.

**FIG. 2.**
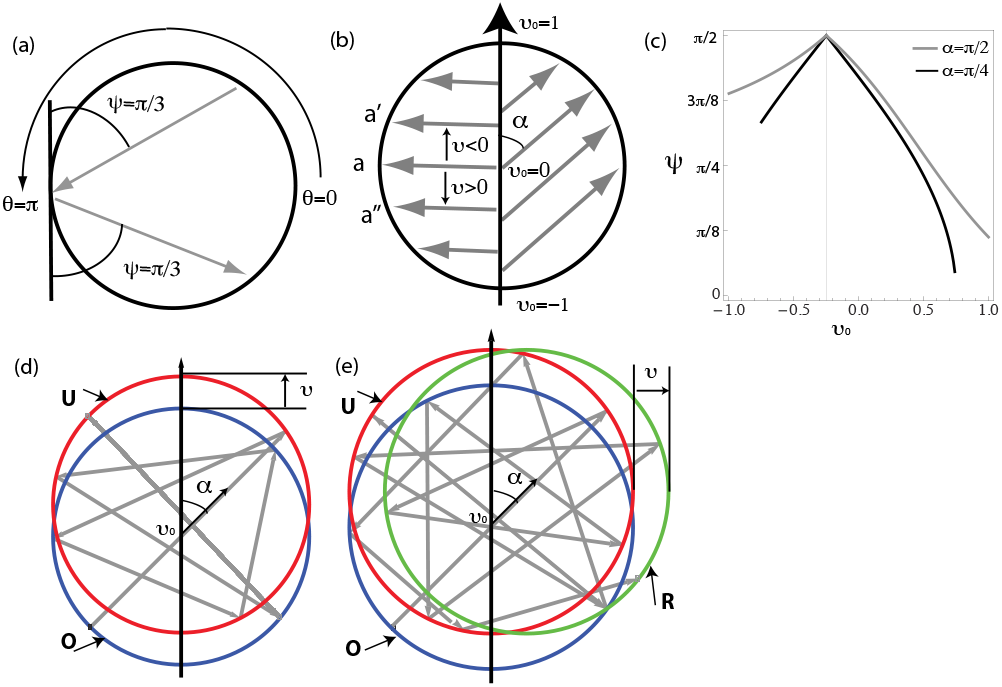
Billiard with moving boundaries: (a) configuration of the system: reflection angle *φ* and collision point *θ*; (b) shifted circle: distance of the shift *v*, angle *α* and coordinate *v*_0_; (c) reflection angle after shift *v* = 0.25; (d) trajectory of the particle for up-down shifting of the circle; (e) trajectory of the particle for up-right-left-down shifting of the circle. In (d) and (e), **O** indicates the original position (blue circle), **U** - the “up” position (red circle), and the left position (green circle).

Next we introduce a new class of billiard - shifting billiard. The circle is shifted in such a way that its center moves distance *v* along the vertical axis, i.e the coordinates of its center is (0, *v*). As the circle boundary is moved, the collision point also changes. For a particle moving along the *x* axis and perpendicular to the vertical axis, i.e. with *θ* = *π* and *φ* = *π*, the collision point becomes *a′* for *v >* 0 (the circle shifted up) or *a″* for *v <* 0 (the circle shifted down), as shown in fig.2 (b). We denote by *α* the angle between the trajectory of the particle and the direction of the circle shift, (*α* ∈ [0, *π*]). We also denote by *v*_0_ the coordinate where the line going through the trajectory of the moving particle intercepts the circle shift directory. Graphical representation of *α* and *v*_0_ is shown in fig. 2 (b). If the angle 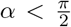, this indicates that the particle moves in the same direction as the circle has been shifted. Otherwise, if the angle 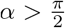, then the particle moves in the direction which is opposite to the direction that the circle has been shifted. Potentially, *v*_0_ ∈ (−∞,∞), but not all *α*-*v*_0_ values will give trajectories contained within the moving circle boundaries. Allowed values of *v*_0_ are within the interval 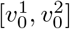, where 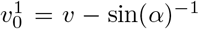 if 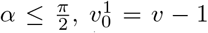, if 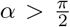; and 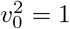 if 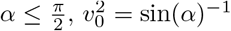 if 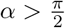.

As the point of the collision changes, so does the angle of the reflection. Figure 2 (c) shows the reflection angle as a function of *v*_0_ for two values 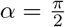 and 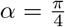 after the circle has been shifted up by *v* = 0.25. For both values of *α*, the reflection angle is equal to 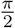 for *v*_0_ = −*v* and decreases for other values of *v*_0_.

We assume an arbitrary periodicity of circle shift corresponding to a time the particle takes to travel from one collision to another. Furthermore, change in position is instantaneous. After the first collision, the circle will return back to its original position with center located at (0,0). Trajectory of the particle moving in the shifting circle are shown in fig. 2 (d). Initial conditions for the particle are *v*_0_ = 0 and 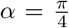, and the circle is shifted by *v* = 0.25. For this particular set of the parameters, the particle is fixed in a period-2 orbit after 6 collisions the with boundary of the circle.

Finally, we introduce a more complex movement of the circle. The circle is shifted according to the following loop: it is moved up (the “up” position, **U**), right (the “right” position, **R**), left (returned to the **U** position), and down (return to the **O** position). Figure 2 (e) shows the trajectory for a particle with the same initial conditions as in 2 (d), only this time it breaks away from period-2 orbit after the circle is shifted to the position **R**.

### B. Model for plating with glass beads

Let Γ denote the unit disk, and *∂*Γ denote it’s boundary. Particle movement within a billiard is defined by the elastic collision rule: the angle of reflection is equal to the angle of incidence [14, 23]. Denote by *q*_*t*_ = (*x*_*t*_, *y*_*t*_) the coordinates of the moving particle at time *t*, and by *ν*_*t*_ = (*u*_*t*_, *w*_*t*_) its velocity vector. Then its position at time *t* + Δ*t* can be computed by

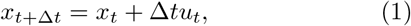

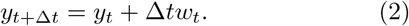

When the particle collides with the boundary *∂*Γ, its velocity vector *ν* gets reflected across the tangent line to *∂*Γ at the point of collision and the new post-collision velocity vector can be computed as

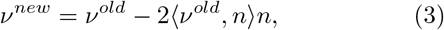

where *n* is the unit normal vector to the boundary and ⟨*ν, n*⟩ denotes the scalar product.

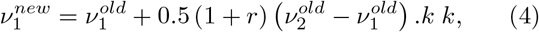

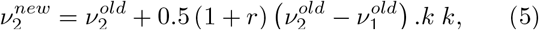

For an arbitrary collision between two particles, the post-collisional velocities are given by:

We have the following set-up for the model: there are *N* glass beads which at a time t are randomly distributed within a unit circle in such a way that they do not overlap, i.e the Euclidean distance between beads centers is large then 2*d*. We assumed that the plating dish is shifted a unit distance (i.e. corresponding to a diameter of the plate) within each step of a loop. All particles are stationary at the start of a simulation, and start moving after collision with the moving boundary *∂*Γ.

We assumed that plating dish is moved up to a unit distance (i.e. corresponding to a diameter of the plate). For an arbitrary time unit *T* = 1, we further assume that the boundary is moved with a velocity *V*. Simulations are run at discrete time intervals with a fixed time step Δ*t* = 0.01 ×*T*. At each time step, we calculate if any particle is close enough to the boundary, or any two particles are close to each other, and create a list of collision events. After each collision, direction and velocity of the involved particle is updated. We assume that velocities of particles are reduced with each time state by a constant, *c*_*v*_, due to the presence of growth media. After running pilot simulations, we found that values *V* = 3 and *c*_*v*_ = 0.99 produced trajectories consistent with the experiment data. During simulations we track position of each particle with respect to the center of the plating dish.

where 0 *≤ r <* 1 is the normal coefficient of restitution, which is typically 0.8 for glass beads [24]; *ν*.*k* denotes the vector product, and *k* is the collision vector, directed from the centre of the second particle to that of the first particle:

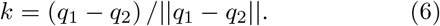

Next, we formulate a stochastic spatially-explicit model of cell dispersal via the movement of glass beads. At time *t* = 0 there are *n* viable cells in suspension which is dispensed onto the middle of the plate. The positions of cells within the plate are modelled using a Gaussian distribution with mean at the center of the plate and variance equal to half of the plate’s radius. Next, we considered two stochastic events: (i) a cell is attached to a bead which is passing within *d/*2 distance from the cell; and (ii) a cell is detached from the bead and is deposited on the surface of the plate. We assume that the probabilities of these events at time *t* are:

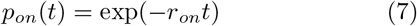

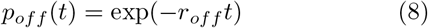

After simulating the dispersal of cells, we calculated which cells were within 1.5 *µm* of other cells, forming clusters. We further assumed that detectable individual CFUs are formed by either separated cells or these clusters.

Parameter inference was performed using Adaptive Multiple Importance Sampling framework [25]. The likelihood was obtained by assuming that the observed number of colonies is Poisson-distributed:

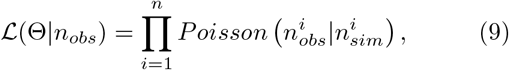

where Θ ={*n, r*_*on*_, *r*_*off*_ *}*; *n*_*obs*_ is the observed number of colonies; *n*_*sim*_ is the number of colonies obtained after simulating the model with parameter set Θ.

## V. RESULTS

First, the plating was carried out with different concentrations of *E*.*coli* cells. We used the “L-shape” movement with 10 glass beads, and either 5 or 10 loops. Results of experiment are shown in Table I. As cell concentration decreased, the mean distance between closest CFUs, *n*_*CFUs*_, and spatial variability of colonies, *CV*_*S*_, increased. The number of colonies was 950-1050 for 10^−5^ dilution, 280-380 for 10^−6^ dilution, and 50-70 for 10^−7^ dilution. It is suggested that for manual counting, the best range of a number of colonies should be between 30 and 300 [26]. Therefore, for further analysis we have chosen 10^−6^ dilution as this gives a good distribution of colonies.

**TABLE I.** Results of the dilution experiment.

| Metrics | 5 loops |  |  | 10 loops |  |  |
| --- | --- | --- | --- | --- | --- | --- |
| | $10^{-5}$ | $10^{-6}$ | $10^{-7}$ | $10^{-5}$ | $10^{-6}$ | $10^{-7}$ |
| $n_{CFUs}$ | 964 | 286 | 59 | 1050 | 382 | 71 |
| $d_{NB}$ | 1.3 | 2.1 | 8.5 | 1.5 | 2.0 | 8.3 |
| $CV_S$ | 0.59 | 0.75 | 1.2 | 0.45 | 0.62 | 0.97 |
| $m_S$ | 7.76 | 3.39 | 1.09 | 8.77 | 3.59 | 1.34 |

Figure 3 displays a summary from the experiments: each row corresponds to a different method of cell plating, number of glass beads and number of loops. We have analysed 40 configurations in total. We observed that the number of CFUs increased when the number of loops increased from 5 to 25. Most of the experiments produced a consistent number of colonies. The exceptions were for cells spread using either 100 loops, or 25 beads and 25 loops, which gave much lower colony numbers. Methods producing the smallest amount of CFUs, also had the largest values of *d*_*NB*_ and *CV*_*S*_.

**FIG. 3.**
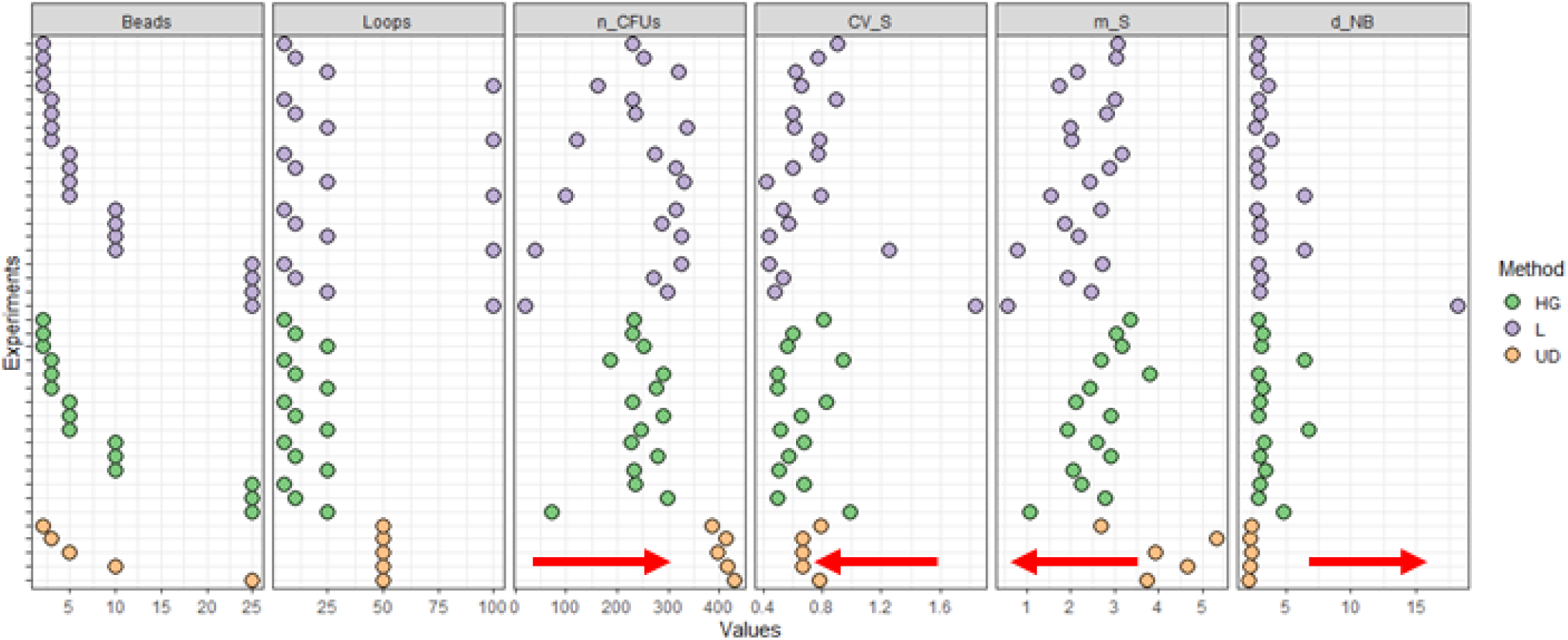
Experimental results. Methods analysed are: hourglass (HG: ↗ ← ↘), L-shape (L: ↑ → ← ↓), and up-down (UD: ↑ ↓). Each row corresponds to a particular experimental setup. The first two columns show the number of beads used and number of loops in each experiment. Red arrows indicate direction from less uniform to more uniform distribution of CFUs.

We could identify three groups of configurations, which produced *n*_*CFUs*_ *>* 300: (i) ‘L-shape’ movement with 2-10 beads and 25 loops; (i) ‘L-shape’ movement with 10-25 beads and 5 loops; and (iii) ‘up-down’ movement with 3-25 beads and 50 loops. However, the latter configuration performed quite poorly with respect to other metrics, especially *d*_*NB*_ and *m*_*S*_. In terms of the number of steps the plate is moved, 50 ‘up-down’ moves are equivalent to 25 ‘L-shape’ moves. Mean distance to the nearest CFUs for the ‘up-down’ method had the smallest values overall. The ‘hourglass’ movement has performed well for 10 loops and 3-25 glass beads in terms of the values of *n*_*CFUs*_, and had one of the lowest values of spatial variability, *CV*_*S*_, however the number of CFUs were not as high as for ‘L-shape’ or ‘up-down’ trajectories. Interestingly, the most natural way to move plates-i.e. wrist movement, was not performing as well as the other methods of plate movement.

Next, we used the model of shifting billiard to investigate the observed spatial patterns in the ‘up-down’ movement. Figure 4 shows experimental images of *E*.*coli* CFUs and density histograms of simulated particle trajectories for the ‘up-down’ movement. For model simulations with *N* beads, we have added an extra bead *N*, but kept initial positions of 1 … (*N* −1) beads the same as in simulations with *N –* 1 beads. It can be seen then even with 50 loops, the trajectories of beads follow the movement of the plate. This mimics exactly how the CFUs are positioned in experiments, i.e. along and parallel to a vertical axis. As the number of beads is increased, there is a possibility of a glass bead escaping the expected trajectory due to a random collision with other glass beads.

**FIG. 4.**
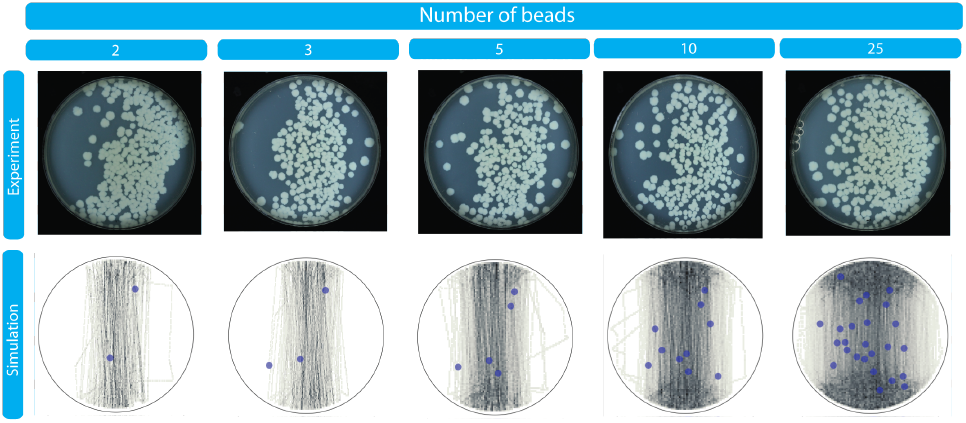
Experimental images of E.coli CFUs and density histograms of simulated particle trajectories for the up-down movement. Columns correspond to a different number of glass beads. Blue circles indicate the initial positions of particles.

Furthermore, we used the model to explain heterogeneities in the observed spatial patterns when using ‘L-shape’ movement. Figure 5 shows representative images of *E*.*coli* CFUs and Figure 6 shows corresponding density histograms of simulated particle trajectories for the ‘L-shape’ movement. It can be seen from the simulations, that when the number of glass beads or a number of loops is low, the surface of the plate can not be fully explored by the moving beads, and particular areas (top left quarter) are hardly visited by any of glass beads at all. In order to improve surface coverage, it is necessary to increase either the number of beads or the number of loops. For example, 5 glass beads with 25 loops, or 10 glass beads and 10 loops, produced a reasonable coverage of the plate surface.

**FIG. 5.**
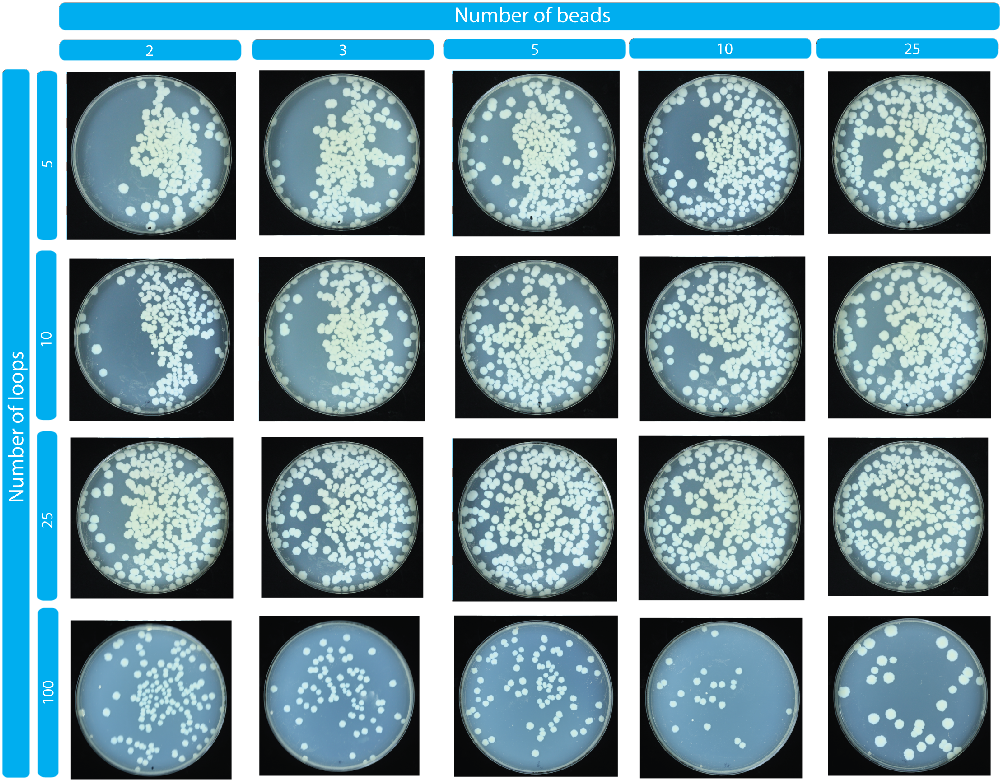
Experimental images of E.coli CFUs for the L-shape movement. Columns correspond to a different number of glass beads, and rows correspond to a different number of loops.

**FIG. 6.**
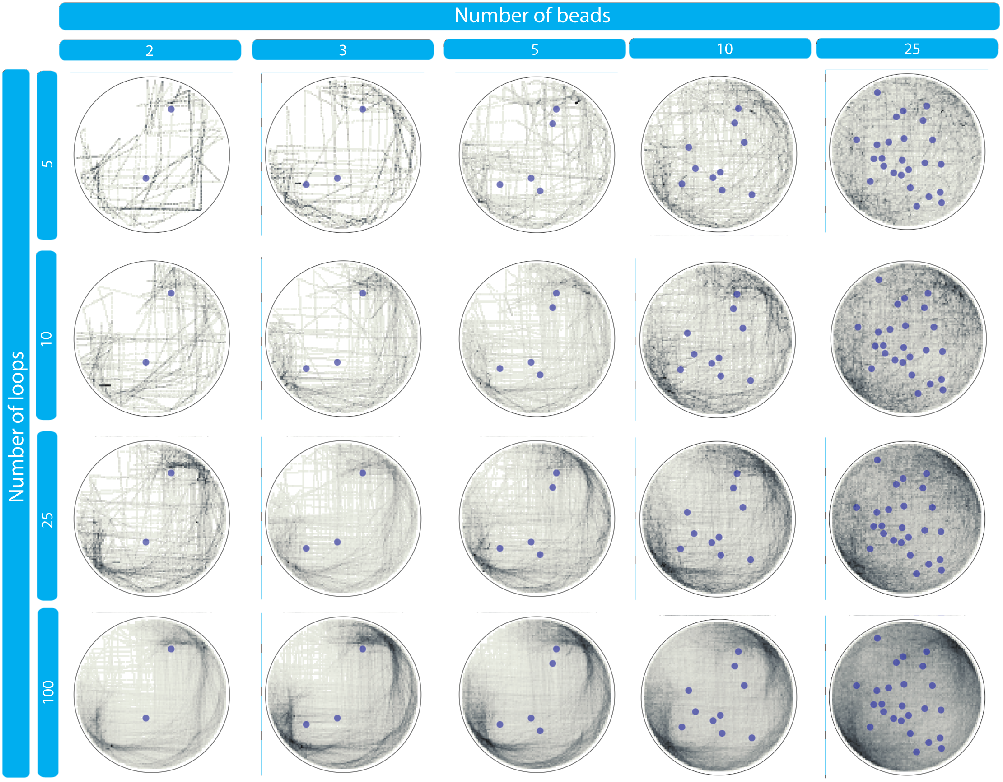
Density histograms of simulated particle trajectories for the L-shape movement. Columns correspond to a different number of glass beads, and rows correspond to a different number of loops. Blue circles indicate the initial positions of particles.

Finally, we have fitted the stochastic model of cell dispersal to experimental data. After pilot investigation, we have fixed *n* = 500. Sampled parameter sets {1*/r*_*on*_, 1*/r*_*off*_} and corresponding log likelihood values are shown in Figure 7(a). We have calculated the maximum likelihood estimate (MLE) as *r*_*on*_ = 0.010416667 and *r*_*off*_ = 0.03571429 (shown as red diamond). Using this parametrized model we simulated cell plating for an ‘L-shape’ trajectory with 2, 5 and 10 beads and the number of loops in the range from 5 to 100 (Figure 7(b)-(d). We have found that 20-30 loops give a maximum number of *n*_*CFUs*_.

**FIG. 7.**
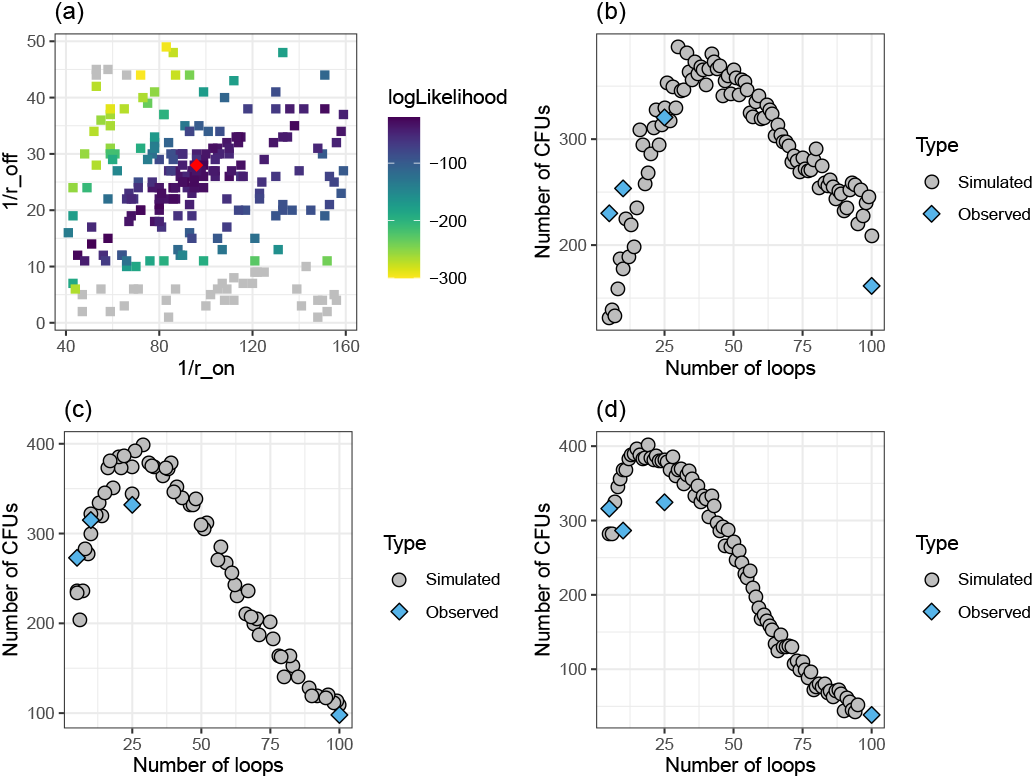
Stochastic model of cell dispersal. Log likelihood values of sampled parameters: gray shows values less than −300; red diamond shows MLE. (b)-(d) Comparison between simulations and experimental data for ‘L-shape’ trajectory with (b) 2 beads, (c) 5 beads, and (d) 10 beads.

**FIG. 8.**
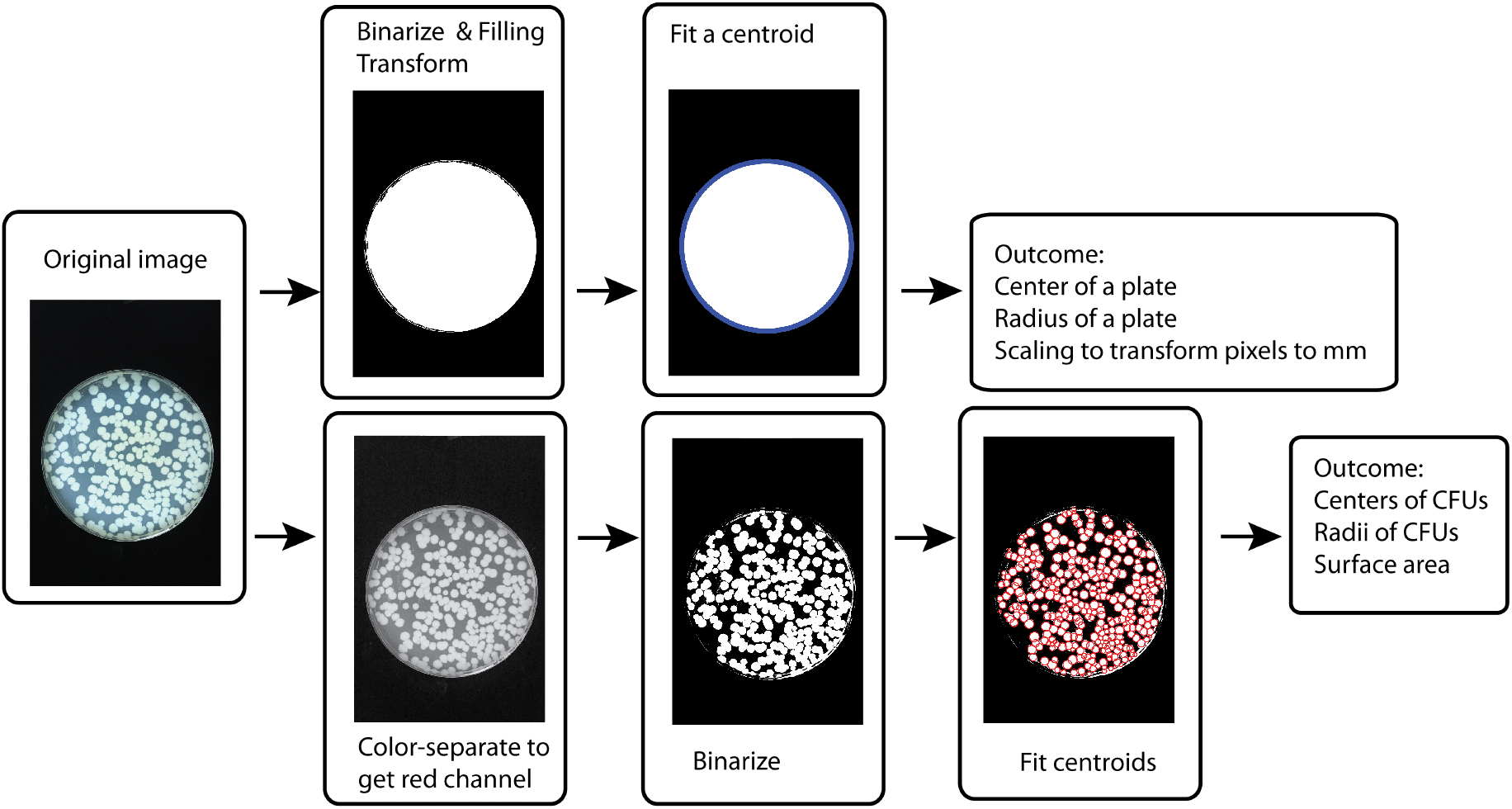
(Supplementary) Flowchart representing image processing. First, image was analysed to detect where plate is. Second, positions of CFUs were detected by fitting centroids.

Our full model agreed with experimental observations of the decrease in the number of colonies when using a very high number of loops. Similar phenomena have been observed when plating central memory cells, which was attributed to cells getting stuck on beads and hence removed during the debeading process [27].

## VI. CONCLUSIONS

In this work we characterize the effectiveness of glass beads for plating cell cultures. We examined the number of colonies, mean distance between closest colonies, and variability and uniformity of spatial distribution of colonies. Our results indicate that the Copacabana method is highly efficient, although care needs to be taken when choosing the trajectory of plate movement, number of beads and number of loops.

An exploratory look at the experimental data revealed some interesting attributes that required further investigation using a mathematical modelling approach. We introduced shifting billiard as a conceptual abstraction of cell plating with glass beads. Numerical analysis of the dynamics of this simple yet efficient billiard indicated a close relationship between plate movement and quality of colony distribution. Simulations show that when the number of glass beads or the number of loops is low, the surface of the plate can not be fully explored by the moving beads. This could be improved by increasing either the number of beads or the number of loops. However, a very high number of loops (i.e. 100) should be avoided, as experimental data and simulations showed that the number of colonies produced was much lower compared to other configurations.

We have analysed the properties of cell spread using digitized images and mathematical modelling. Optical density (OD) readings have been used to evaluate the number of CFU based on the time it takes to reach the predetermined OD of 0.1 when compared with a reference standard curve [28]. However, this technique would not provide the necessary details for the spatial analysis we performed.

In conclusion, we recommend using ‘L-shape’ trajectory with 2-10 beads and 25 loops which should produce good colony separation in terms of the number and spatial spread of colonies. Recently, automated systems have been utilised for high-throughput cell handling and analysis [29]. Manual transformation and spreading on agar plates are still in otherwise fully automated liquid handling platforms [30]. Therefore, we believe that our results should be of interest when using automated platforms and could guide programming of automatic cell plating protocols.

## Appendix A

### Image analysis

Each image has been binarized with a low threshold in order to find the boundaries of a plate; this gives scaling for transforming pixels into millimeters. Next, we extracted single-channel images corresponding to each of the color channels in image. We have found that working with image in red channel improved detection of CFUs. Finally, we have fitted centroids representing CFUs by varying a threshold of binarization.

